# COMT Val158Met polymorphism associated with greater susceptibility to framing effects in healthy older adults

**DOI:** 10.1101/790808

**Authors:** Carson M. Quinn, Mia Borzello, Ali Zahir, Joel H. Kramer, Winston Chiong

## Abstract

Age-related neural changes may compromise older adults’ decision-making, increasing their risk of fraud and financial abuse. One manifestation of nonrational influences on decision-making is susceptibility to “framing effects,” in which decisions are biased by irrelevant contextual features of how choice information is presented. We investigated whether polymorphisms in genes related to dopamine neurotransmission (COMT) and neurodegeneration (ApoE) influence the susceptibility of older adults to framing effects. We administered an online test of susceptibility to framing effects to a cohort of 113 healthy older adults who had undergone genetic testing for COMT and ApoE genotype. The task required the participant to choose a risky or safe option in pairs of situations that were monetarily equivalent but differed in whether the choice was framed in terms of gains or losses. A general linear model was used to test for associations between inconsistency in choice across the set of choice pairs and these genotypes, controlling for age, education, gender and traditional measures of executive function. While no association to framing effects was found for ApoE, the Valine allele of COMT Val158Met was significantly associated with greater susceptibility to framing, although the association was no longer significant after adjustment for demographic covariates. Our results suggest that greater frontal dopamine concentrations associated with the COMT Met allele are protective against less consistent decision making in older adults. When compared to previous findings in young adults, our findings provide additional support for an inverted-U shaped model of prefrontal dopamine function.

## 1. Introduction

Given population aging and other global sociodemographic trends, older adults are now expected to make more consequential decisions about their health and finances at more advanced ages than ever before. These decisions have important implications for their families and for society as a whole; in 2013, 36% of U.S. household wealth was managed by adults 65 and over (US Census Data). Unfortunately, age-related cognitive changes may compromise many older adults’ decision-making, placing them at risk for significant losses and financial abuse, which is the most common form of elder abuse (Aciemo et al., 2010). For example, using real-world population-level data Agarwal and colleagues have demonstrated less advantageous decision-making in advanced age across several consumer financial behaviors (Agarwal et al., 2009).

While classical theories of decision-making have focused on analytic processes, empirical behavioral research suggests that decisions are often guided by more heuristic or intuitive modes of decision-making, especially under conditions of uncertainty (Gilovich et al, 2002). This process of decision-making is characterized by urgency and lack of premeditation, and underlies decisions that are influenced by impulsivity. Such heuristic decision-making is manifested in “framing effects” in which decisions are biased by irrelevant contextual factors in how information is presented; as when someone would be willing to take a medication with an 80% chance of not having a severe side effect, but unwilling to take a medication with a 20% chance of having a severe side effect (Tversky and Kahneman, 1981). Recently, functional neuroimaging studies have associated framing effects with activity in the prefrontal cortex, anterior cingulate cortex, and limbic regions (De Martino et al., 2006; Xu et al., 2013). These brain regions are primary targets of midbrain dopaminergic systems known to be significantly affected in cognitive aging, with declines of nearly 10% per decade reported in dopamine cell number (Ma et al., 1999) and postsynaptic receptor density (Kaasinen et al.,2000; Inoue et al.,2001). As such, these systems are promising targets of inquiry into neurobiological processes underlying decision-making in aging.

For example, the catechol-*O*-methyltransferase (COMT) enzyme is the primary route for dopamine degradation in prefrontal cortex (Gogos et al.,1998), and enzyme activity strongly influences synaptic dopamine concentration in prefrontal cortex (where the dopamine transporter has low expression) (Lewis et al.,2001). The rs4680 functional polymorphism in the COMT gene results in a valine (Val) to methionine (Met) substitution at codon 158, with the Met allele enzyme having 40% less activity than the Val allele enzyme (Lotta et al.,1995), which is believed to result in higher prefrontal synaptic dopamine concentrations in Met/Met than Val/Val individuals. Numerous studies have probed the role of the COMT Val158Met polymorphism in behavior and cognition, with the Met allele being associated with stronger executive function (Wishart et al., 2011) and neuroticism (Enoch et al.,2008). However, the complexity of the gene-environment interaction is apparent in the numerous studies that have found conflicting or unclear results (Tsai et al.,2003; Smyrnis et al.,2007). One prior study examining COMT and framing effects in young adults reported a modest, but statistically significant effect of the Met allele, resulting in greater susceptibility to framing effects (Gao et al.,2016). Interestingly, the prevailing hypothesis on prefrontal availability posits an inverted-U-shaped relationship between dopamine tone and prefrontal cognitive function, with both high and low dopamine levels being less advantageous than a moderate level (Witte and Floel, 2012). This relationship has been supported both by studies of cognitive performance with dopamine-enhancing interventions (Cai and Arnsten, 1997; Mattay et al., 2003) as well as in studies of neuropsychiatric conditions (Williams-Gray et al.,2008; Na et al.,2018). By this hypothesis, there is reason to suspect that the influence of COMT Val158Met on susceptibility to framing effects might differ in an older population with decreased dopamine signaling, as compared to a younger population. In this study, we investigated the relationships among aging, the heuristic versus algorithmic decision-making differences manifested in framing effects, and the COMT Val158Met polymorphism. We administered an online task measuring framing effects to 88 healthy participants aged 62-92 who had been genotyped at the COMT 158 locus, and who had also been genotyped for the ApoE gene, a risk factor for Alzheimer’s disease. We hypothesized that in this population, the Val allele would be associated with greater susceptibility to framing effects.

## 2. Methods

### 2.1. Study participants

A sample of 113 neurologically healthy, community dwelling older adult participants was included from a cohort at the University of California, San Francisco Memory and Aging Center that includes in-person measurements (such as standardized cognitive testing, neurological examination, and biomarkers) as well as online web-based behavioral assessments completed at home. Participants were between 62 and 92 years of age and were enrolled as neurologically normal on the basis of informant interview, neurological examination, and cognitive testing. All subjects were reviewed after in-person testing at a consensus case conference with a neuropsychologist and neurologist; exclusion criteria included a syndromic diagnosis of dementia or mild cognitive impairment, a Mini-Mental State Examination score less than 26, severe psychiatric illness (e.g., schizophrenia, bipolar disorder), neurological condition that could affect cognition (e.g., epilepsy, Parkinson’s disease), substance use diagnosis in the last 20 years, significant systemic medical illnesses, current depression (Geriatric Depression Scale ≥15 of 30), or subject or informant report of significant cognitive decline over the previous year. All participants provided written informed consent, and the study protocol was approved by the UCSF Committee on Human Research.

### 2.2. Genotyping

DNA was extracted from saliva samples obtained via cheek swab using standard procedures. Subjects were genotyped for the three Val158Met COMT polymorphisms (Met/Met, Val/Met, Val/Val) and ApoE. COMT (rs4680) genotypes were obtained using a commercially available TaqMan® SNP genotyping assay (C_25746809_50) from ThermoFisher on a LightCycler® 480 System using standard end-point genotyping protocol provided by the manufacturer. ApoE genotype was determined with a TaqMan Allelic Discrimination Assay for the two SNPs, rs429358 and rs7412, on an ABI 7900HT Fast Real-Time PCR system (Applied Biosystems, Foster City, California) using the manufacturer’s instructions. The AccuPrime™ Taq High Fidelity DNA Polymerase kit was used for the polymerase chain reaction. The reaction contained a 6-FAM labeled forward primer (/56-FAM/GCGACTACGTGGTCTACTCG) and a reverse primer (AGGACCCTCATGGCCTTG) (Primers referenced from Lichter et al HMG 1993). The PCR reactions were then sent for fragment analysis on an AB3730XL with a LIZ1200 size standard. This method is a modification of the gel-based assay set forth by (Chen et al., 1999).

### 2.3. Framing effects task

Participants performed the behavioral experiment online through a web-based instrument. The survey instrument was designed and implemented using the online survey platform Qualtrics© (<https://www.qualtrics.com/>). Each participant received an individualized e-mail containing a unique web link to complete the task; this link could be used to return to the task if it was not completed in one sitting. Compensation for task completion was e-mailed to participants as an electronic gift card.

The behavioral task was adapted from an in-person computer-based task that has been previously described (De Martino et al., 2006). Participants were presented with a total of 74 trials. Each trial consisted of a cue screen, indicating one of four different starting endowments: $2, $5, $10, or $20 (e.g. “You receive $10”). On the next screen, participants were asked to make a choice between two options- a sure option and a gamble option. The sure option was framed as either the amount of money kept from the initial amount (“Gain” frame; e.g., “Keep $2” out of initial endowment of $10) or as the amount lost from that initial amount (“Loss” frame; e.g., “Lose $8” out of an initial endowment of $10—note that this is numerically equivalent). In both the Gain and Loss frames, the gamble option was consistently depicted as a pie chart showing the probability of winning or losing all of the starting endowment in the given trial. In the 64 trials of interest, four different probabilities were used: 20, 40, 60, 80%. These four probabilities were counterbalanced with the four starting amounts across frames. The expected monetary values of the options were balanced in each trial of interest; for example, in one trial with a starting endowment of $10, participants were asked to choose between a sure option “Keep $2” and a gamble option with a 20% chance of keeping all $10 and an 80% chance of losing all $10.

Given concern in the field of behavioral research that online tasks might not foster adequate motivation to understand the task, it is typical to regularly assess participant understanding. To monitor for the active engagement and understanding of participants during the task, we designed the task with interspersed “catch trials” in which the expected monetary values were highly unbalanced, and in which one option was clearly preferable to the other (given conservative assumptions about participants’ individual risk aversion). Across both frames of the catch trials, half involved gambles that were markedly preferable to the sure option (i.e., a gamble with 95% probability of winning the entire starting endowment versus a sure option of 50% of the starting endowment), and half involved gambles that were markedly inferior to the sure option (i.e., a gamble with 5% probability of winning the entire starting endowment versus a sure option of 50% of the starting endowment). Following prior studies, in our initial design we defined adequate engagement and understanding of the task as correctly answering at least 7 of the 10 catch trials (De Martino et al., 2006). In order to avoid inclusion of data from participants who misunderstood or lost focus on the task, our analyses excluded participants who did not meet this threshold.

### 2.4. Data analysis

The primary outcome of interest was susceptibility to framing effects, quantified as the difference between the percentage of trials in which a participant chose to gamble in the loss frame and the percentage of trials in which that participant chose to gamble in the gain frame. Because the set of choices presented in the gain frame was numerically equivalent to the set of choices presented in the loss frame, a difference in the willingness to gamble reflects the effect of framing on a given participant.

For each gene of interest, we constructed a general linear model with the difference in gambling frequency from the loss to the gain frame as the outcome variable, and gene status, age, years of education, and gender as predictor variables. For the COMT locus, gene status was quantified by the number of valine alleles. For the ApoE locus, gene status was entered as a risk scale increasing with risk; with E2/E2 as 1, followed sequentially by E2/E3, E3/E3, E2/E4, E3/E4, and finally E4/4 as 6. Change in gambling frequency was modelled with a Gaussian distribution and identity link. To evaluate for contributions of traditional measures of executive function to susceptibility to framing effects, a secondary analysis was performed in a subset of participants who had undergone in-person standardized cognitive testing within 18 months of online behavioral task performance. This included the same predictors as the original model, including a further predictor variable representing a summary of executive function. This summary executive function measure was computed as the average of z-scores for backwards digit span (a test of attention), Stroop interference (a test of response inhibition), a modified Trails test (a test of set shifting), semantic fluency, and design fluency.

In a supplementary analysis, we assessed whether some groups were systematically excluded from the primary analysis due to our use of catch trials. We constructed a linear regression model with the binary variable of exclusion based on catch trial performance as the outcome variable, and gene status, age, and executive function as predictor variables in successive analyses. All statistical analyses were conducted in Stata 14.2 (StataCorp, College Station, TX); a two-tailed P-value of <0.05 was considered significant.

## 3. Results

### 3.1. Cohort characteristics

Demographic information, score on the bedside executive function battery, and the primary outcome measure of the framing effects bias are presented in Table 1 for the full sample, as well as within COMT and ApoE genotype subgroups. Age, education, and executive function were similar between the genetic groups. As indicated in the table, 35% of participants did not meet our performance threshold on catch trials, leaving 73 of the original 113 participants with a measure of susceptibility to framing effects. The COMT Val158Met polymorphism was in Hardy-Weinberg equilibrium in our population (chi-square = 0.012, p = 0.914), with 26 Met/Met, 57 Met/Val, and 30 Val/Val individuals. The ApoE polymorphism was in Hardy-Weinberg equilibrium in our population (chi-square = 0.416, p = 0.937), with 0 E2/E2, 10 E2/E3, 2 E2/E4, 70 E3/E3, 24 E3/E4, and 2 E4/E4 individuals.

**Table 1.** Cohort characteristics of the full sample, and divided by COMT genotype (left) and APOE genotype (right).

|  | **Full sample** | **Met/Met** | **Met/Val** | **Val/Val** | **Not E4 Carrier** | **E4 Carrier** |
| --- | --- | --- | --- | --- | --- | --- |
| *N* | 113 | 26 | 57 | 30 | 80 | 28 |
| *Age* | 76.3<br>(5.8) | 77.4<br>(5.6) | 76.3<br>(5.4) | 75.2<br>(5.3) | 76.1<br>(5.8) | 76.9<br>(5.9) |
| *Education years* | 17.6<br>(2.1) | 17.2<br>(2.3) | 17.5<br>(2.2) | 18.3<br>(1.6) | 17.5<br>(2.1) | 17.5<br>(2.2) |
| *% Female* | 59.3 | 50 | 59.6 | 66.7 | 60 | 53.6 |
| *Executive Function score* | -.054<br>(.486) | -.009<br>(.467) | -.036<br>(.495) | -.125<br>(.496) | -.088<br>(.511) | .098<br>(.410) |
| *N who completed Framing effects* | 73 | 17 | 35 | 21 | 47 | 23 |
| *Change in gamble frequency percent* | 19.2<br>(19.2) | 12.7<br>(11.9) | 18.5<br>(18.1) | 25.6<br>(23.9) | 20.9<br>(20.8) | 17.7<br>(15.7) |

Unless otherwise specified, values are listed as means with standard deviations.

### 3.2. Framing effects task

As in previous studies utilizing versions of the framing effects task, participants were more likely to gamble in the loss frame than in the gain frame (p<.001); gambles were chosen in 30.7% (95% CI= 25.9-35.4) of trials of interest in the gain frame, compared with 49.8% (95% CI =44.6-55.1) in the loss frame (Fig 2).

**Fig 1.**
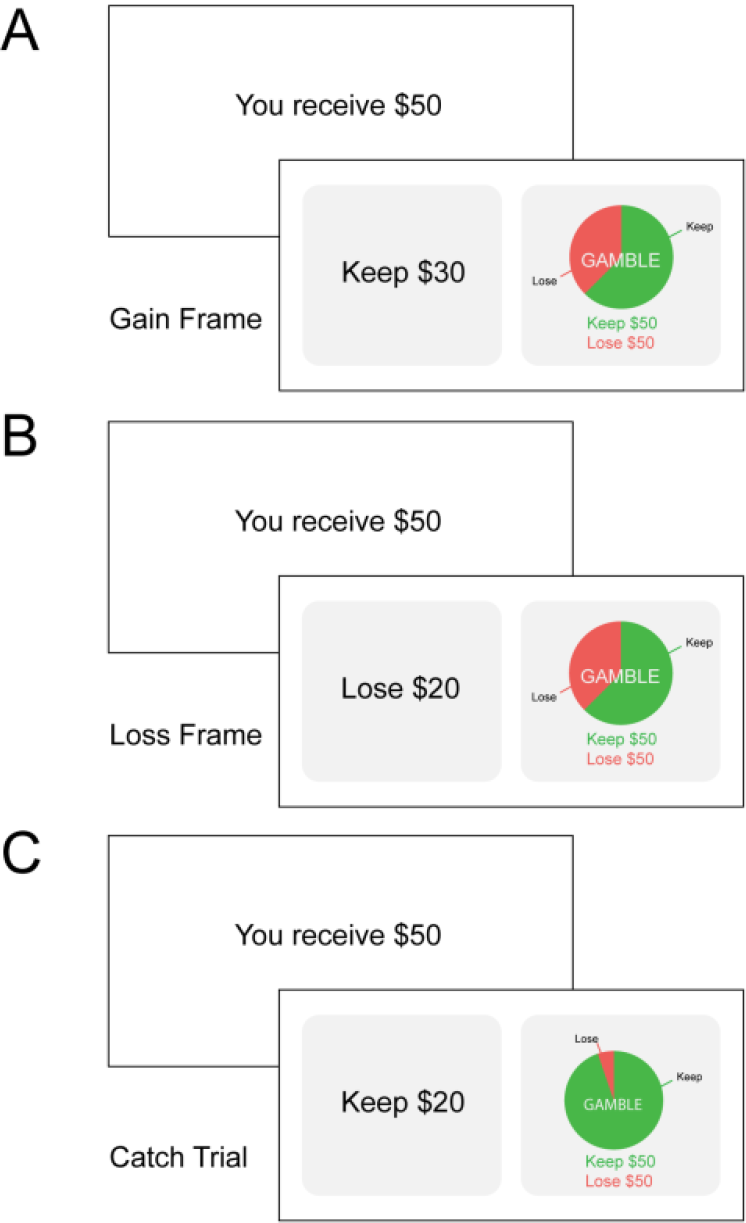
Experimental Task. Each trial consisted of a cue phase and a decision phase. In the cue phase, participants are presented with a screen showing a starting amount or endowment. Then, in the decision phase, they are asked to choose between the safe option and the gamble option. The safe option was presented in two different conditions: (A) a gain frame (“Keep”) and (B) a loss frame (“Lose”). The gamble option was presented the same way, regardless of frame.

**Fig 2.**
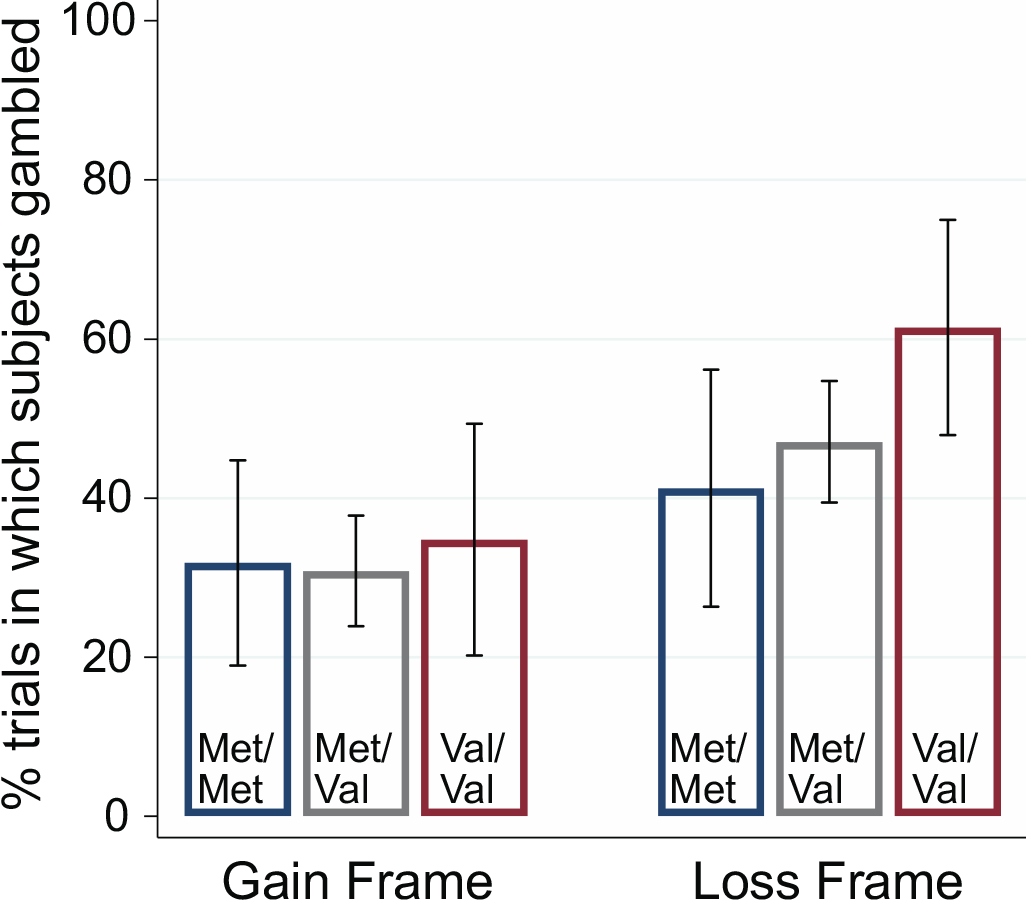
Differences in gambling frequency between Loss and Gain Frame. Participants of all genotypes chose the gamble option more frequently in the loss frame than the gain frame. The same relationship held for each individual genotype, with the Valine allele showing a dose-dependent relationship to gambling frequency in the Loss Frame. Error bars signify 95% confidence intervals

ApoE genotype was not significantly associated with susceptibility to framing effects, either alone, or when controlling for demographics and executive function (p=0.457).

The COMT Val158Met polymorphism was significantly associated with susceptibility to framing effects; the effect of the COMT Val_158_ allele load was 6.5 (95%CI 0.520 – 12.5, p=0.03) This can be interpreted to mean that each Val_158_ allele is associated with an 6.5% increase in the difference in gambling frequency between the gain and loss frames. However, when controlling for the prespecified covariates age, gender, and education, the association no longer met the threshold for statistical significance and the effect of COMT Val_158_ allele load was 6.0 (95% CI - 0.412 – 12.4, p=0.067). Age, gender and educational attainment were not independently associated with susceptibility to framing effects, and when each covariate was sequentially included in the model without the other covariates, a statistically significant association between framing effects susceptibility and COMT was found. In 67 participants who also underwent standardized cognitive testing within 18 months of performing the framing effects task, the association maintained statistical non-significance after also controlling for a summary measure of executive function, with an unchanged coefficient of 6.0 (95%CI - 0.454 – 12.4, p=0.068); executive function was not independently associated with susceptibility to framing effects (p=0.691). This increasing discrepancy in gambling frequency with Val_158_ allele load was attributable to dose-dependent increases in gambling frequency in the loss frame, while gambling frequency in the gain frame was consistent across genotypes. (Fig 2)

### 3.3. Supplementary analyses

Neither the APOE genotype (p=0.184) or COMT Val158Met genotype (p=0.911) was significantly associated with exclusion from analysis for poor performance on catch trials.

## 4. Discussion

In this study, we investigated the relationship between the COMT Val158Met polymorphism (and the ApoE polymorphism) and susceptibility to framing effects in older adults between 62 and 92 years of age. While ApoE genotype was not associated with susceptibility to framing, we found a dose-dependent effect of COMT Val_158_ allele load with increasing susceptibility to framing effects. This effect was attributable to increased gambling frequency in the loss frame. Within our sample of older adults age, gender, educational attainment and executive function were not associated with susceptibility to framing effects. However, once our model was adjusted for these prespecified demographic covariates, the association no longer achieved significance. We therefore cannot exclude the alternate hypothesis that the association between susceptibility to framing effects and COMT genotype is a result of demographic confounding from the combined effect of age, gender, and education. Further study is necessary to better determine whether the association reflects a real causal relationship or is simply a result of confounding.

Our findings contrast with work in college-age adults reported by Gao and colleagues (2016), in which the COMT Met_158_ allele was associated with increased susceptibility to framing effects, which in that study was driven by increased gambling frequency in Met carriers in the loss frame. This discordance in findings across different age cohorts parallels those reported in delay discounting. In adolescents, more impulsive choice (i.e., greater discounting of delayed rewards) is associated with the Met allele (Paloyelis et al.,2010; Smith and Boettiger, 2012); while in young-to-middle-aged adults, more impulsive choice is associated with the Val allele (Smith and Boettiger, 2012; Boettiger et al., 2007).

Both sets of findings can be reconciled within an inverted-U-shaped model of prefrontal dopamine function, in which prefrontal dopaminergic circuits are responsible for regulation of intertemporal choice and of consistency across different frames of the same choice (Fig 3). According to this model, in younger cohorts with greater baseline dopaminergic activity, increased prefrontal dopamine availability due to reduced enzymatic degradation with the Met allele results in reduced prefrontal function; and thus, increased impulsivity and susceptibility to framing. However, in older cohorts after age-related reductions in dopaminergic activity, increased prefrontal dopamine availability associated with the Met allele results in increased prefrontal function; and thus, decreased impulsivity and susceptibility to framing.

**Fig 3.**
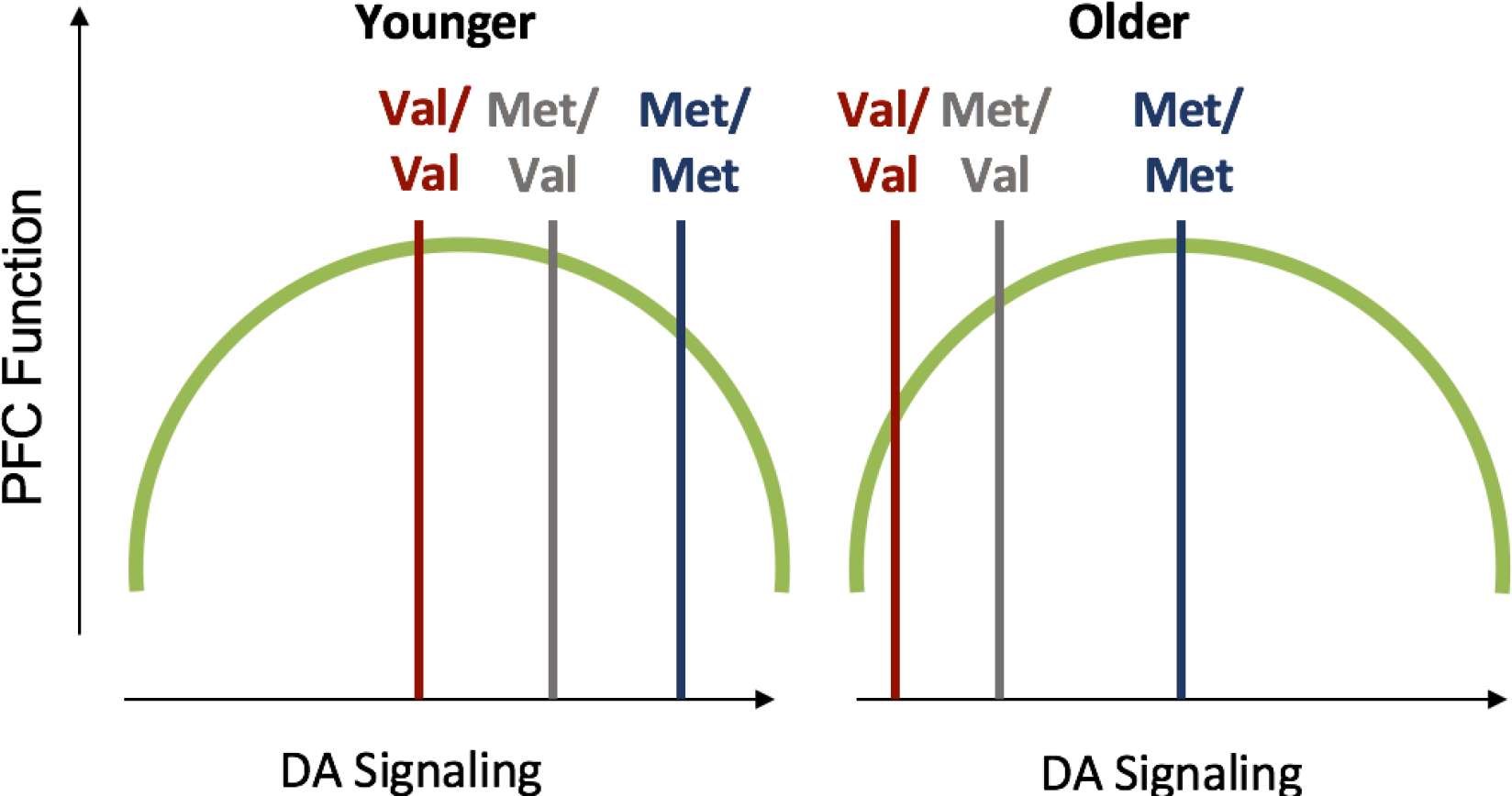
Inverted U-shaped dopamine response model. The left curve shows model-predicted levels of dopamine (DA) signaling for the three genotypes in young individuals, while the right curve shows the same genotypes in older individuals. By this theoretical model, Val/Val individuals have optimal prefrontal cortex (PFC) function early in life when there is greater DA signaling at baseline, but Met/Met individuals show more optimal PFC function in later life once DA signaling has decreased.

### 4.1. Limitations

The principal limitation of our study was its small sample size, exacerbated by the fact that only 73 of the 113 participants recruited for the study performed adequately enough on “catch trials” to be included in the analysis. The online framing effects task was both long and cognitively demanding, and similar studies have also reported a high proportion of participants excluded for poor catch trial performance indicating low engagement with the task (Gao et al., 2017). In this case, while online task administration allowed for data collection that was more convenient to participants, many participants may have been less engaged during off-site testing than if they had performed the task in a controlled testing environment. As a result our sample size was smaller than expected, limiting our ability to conclude whether the association found between susceptibility to framing effects and this COMT polymorphism was real or a result of demographic confounding. Further research is needed to better characterize the relationship. It is also notable that our cohort was highly educated, which might diminish generalizability to other populations.

### 4.2. Conclusions

In conclusion, our study suggests an association between the Val allele of COMT Val158Met and greater susceptibility to framing effects in older adults, but we cannot exclude the alternate hypothesis that the association results from demographic confounding. This suggests that in aging, decreased frontal dopamine concentrations predispose individuals to more impulsive, less rational decision-making. Given our small sample size, further study is necessary to determine the true nature of the relationship between COMT and decision making in older adults.

## Acknowledgments

The authors thank our research participants at the UCSF Memory and Aging center for their gracious contributions to our research. We also thank Rowan Saloner and Alexander Beagle for assistance in participant recruitment and task administration; Anna Karydas, Ryan Fitch and Giovanni Coppola with assistance on genotyping and sample handling; and Ming Hsu and Andrew Kayser for valuable discussion.

